# Chemogenetic Inhibition of the Ventrolateral Orbitofrontal Cortex Disrupts Novelty-Guided Updating of Spatial Memories and Alters Hippocampal Engram Reactivation

**DOI:** 10.64898/2026.07.16.738971

**Authors:** K Kim, D Yu, WF Wade, R Zdziarski-West, Z Ahmad, M Cutler, G Romanelli, A Daki, SL Grella

**Affiliations:** Loyola University Chicago, Department of Psychology (Neuroscience Program), Chicago, IL, USA, 60660; Department of Neuroscience, Feinberg School of Medicine, Northwestern University, Chicago IL, USA, 60611

**Keywords:** Orbitofrontal Cortex, Memory-Updating, Prediction Error, Engram, Novelty-Detection, Dentate Gyrus, DREADDs

## Abstract

**Rationale:** Adaptive behavior requires revising memories when outcomes deviate from expectations. The orbitofrontal cortex (OFC) is implicated in representing outcome expectancies, but its role in memory-updating, beyond value-based learning, remains unclear.

**Objectives:** We tested whether the ventrolateral OFC (VLO) contributes to hippocampal-dependent memory-updating in the Objects in Updated Locations (OUL) task, where novelty-driven exploration provides a behavioral readout consistent with successful updating of spatial information

**Methods:** Using inhibitory DREADDs, in male and female (Swiss Webster and C57BL/6) mice, we suppressed VLO activity during the Updating session, when new object-location information was introduced.

**Results:** Control mice preferentially explored the updated object-location configuration during Updating and discriminated between updated and novel object-location configurations at Test, whereas VLO-inhibited mice did not. VLO inhibition when no updating demand was present did not alter exploration of novel object-location configurations or memory for the original spatial configuration at Test.

**Conclusions:** These results indicate that VLO activity is selectively required when novelty must be interpreted relative to prior experience, implicating this region in evaluating change rather than detecting it. To assess hippocampal ensemble dynamics, we tagged neuronal ensembles, in the dorsal dentate gyrus (dDG) recruited during Updating, followed by quantification of overlap between Updating-tagged cells and ensembles reactivated at Test. We found that VLO inhibition reduced reactivation of Updating-tagged dDG ensembles, paralleling the behavioral impairments observed in the OUL task. These findings support a role for the VLO in hippocampal-dependent memory-updating and extend OFC models beyond reward contexts to include predictive updating of spatial memory representations.

**Significance Statement:** Using a spatial paradigm that distinguishes detection of changed information from incorporation of that information into an updated hippocampal representation, we found that ventrolateral orbitofrontal cortex (VLO) activity is required for memory-updating when new information is present. We found behavioral deficits accompanied by impairments in hippocampal ensemble representations, indicating VLO activity is necessary for memory-updating at the hippocampal engram level. Through this mechanism, the OFC supports the flexible updating of expected outcomes when environmental conditions shift, allowing for the re-evaluation of previously learned associations. These findings extend models of orbitofrontal cortex function beyond reward-guided behavior and identify a cortical contribution to predictive spatial memory processes. Disruption of this computation may contribute to the cognitive rigidity observed in neuropsychiatric disorders.

## 1. Introduction

Memory-updating is thought to occur when retrieved representations are rendered labile by prediction errors, allowing new information to be incorporated into existing memory traces (Exton-McGuinness et al., 2015; Groves et al., 2025; Milton et al., 2023; Nader et al., 2000; Wahlheim & Zacks, 2025). Adaptive behavior in dynamic environments depends on the brain’s capacity to update memories when expectations are violated (Lourenco & Casey, 2013). The orbitofrontal cortex (OFC) plays a critical role in assigning salience to outcomes, predicting and representing expected rewards and their associated values (Liu et al., 2025; Rolls et al., 2020), and incorporating new information when predictions are no longer valid (Sakaki et al., 2011; Takahashi et al., 2009). Although traditionally examined in the context of value-based decision making (Murray et al., 2015; Qiu et al., 2024; Zhou et al., 2021), emerging evidence indicates that the OFC also contributes to memory-updating by signaling reward predictions, guiding representational change, and interacting with mnemonic systems (Costa et al., 2023; Mızrak et al., 2021; Namboodiri et al., 2019; O’Neill & Schultz, 2013; Stalnaker et al., 2018; Tsukano et al., 2026; Witkowski et al., 2025). Mechanistically then, the OFC supports flexible updating of expected outcomes when environmental conditions shift, allowing for re-evaluation of previously learned associations (Aguirre et al., 2024; Gardner et al., 2017, 2020; Schiereck et al., 2026). Understanding how the OFC supports these memory dynamics is essential for elucidating the neural mechanisms underlying cognitive flexibility and its disruption in neuropsychiatric disorders.

The ventrolateral OFC (VLO) projects to many subcortical structures (Cavada et al., 2000; Reiten et al., 2023), is linked to behavioral flexibility (Butkovich et al., 2025) and integrates sensory, reward, and associative information (Rolls et al., 2020; Rolls & Grabenhorst, 2008). VLO inactivation impaired flexible behavior by disrupting the ability to integrate outcome expectancies (Sarlitto et al., 2018). Thus, it is anatomically and functionally poised to alter episodic representations. However, the causal contribution of the VLO to memory-updating remains unclear. Given its putative role in integrating associative and mnemonic information, we tested whether neural ensembles within this region are necessary for a hippocampal-dependent spatial memory-updating task (Exps. 1 & 2) and whether its inactivation disrupts updating of hippocampal representations (Exp 2).

The Objects in Updated Locations (OUL) Paradigm (Kwapis et al., 2020), can be used to examine hippocampal-dependent spatial memory-updating. In this task, mice first learn the locations of several identical objects during Training. During Updating, one object is moved to a new location, violating the animal’s previously learned spatial expectations. At Test, mice are exposed to all object-locations, including the updated location as well as a novel location. Because mice naturally prefer novel spatial configurations, increased exploration of the updated location is interpreted as reflecting successful incorporation of new spatial information into an existing memory representation. In short, the OUL task was developed to preferentially assess memory-updating by requiring animals to reconcile previously learned and newly encountered spatial information.

Here, we employed inhibitory DREADDs (Designer Receptors Exclusively Activated by Designer Drugs) (Roth, 2016) to selectively suppress activity in the VLO during the memory-updating session, enabling a causal test of its role in this process (Exp 1). We hypothesized that disrupting VLO activity would impair memory-updating, thereby illuminating how OFC circuitry contributes to cognitive flexibility and how its dysfunction may underlie maladaptive rigidity seen in certain disorders (Giommi et al., 2023; Ottaviani et al., 2016; Waltz, 2017). To further investigate whether VLO inhibition altered hippocampal representations associated with memory-updating, we replicated this experiment while tagging neuronal ensembles in the dorsal dentate gyrus (dDG) using a viral TetTag system (Doucette et al., 2020; Grella et al., 2022; Reijmers et al., 2007). Neurons recruited during the Updating session were tagged and their reactivation at Test was subsequently quantified (Exp. 2). We selected the dDG because of its established role in contextual memory-updating, pattern separation, and hippocampal engram formation (Doucette et al., 2020; Grella et al., 2019, 2020, 2022; Groves et al., 2025; Rolls, 2013; Schmidt et al., 2012; Yassa & Stark, 2011). The OUL task requires animals to distinguish between highly similar spatial representations that differ only by a single object-location change, a computation thought to rely heavily on DG processing. We therefore tested whether VLO inhibition altered the reactivation of updating-related ensembles within this hippocampal subregion.

## 2. Materials and Methods

### 2.1 Animals

All experimental procedures were conducted in accordance with protocols 3509 and 3513 approved by the Institutional Animal Care and Use Committee (IACUC) at Loyola University Chicago. A total of 64 mice (Swiss Webster mice: Exp 1: n=40, 20M, 20F) and 24 inbred (C57BL/6 mice: Exp 2: n=24, 12M, 12F) were included in this study. Mice were obtained from Charles River Labs and subsequently bred in-house. They were approximately 35 days old at the start of the experiment, weighing between 20-35g. Mice were housed in groups of 2-3 per cage and kept in a temperature and humidity-controlled colony room (temperature: 73°F, humidity: 58%) on a regular light cycle (12:12h light ON/OFF, lights ON at 6:00am). Experiments were run during the light part of the cycle under red light to eliminate glare in the testing chamber. Cages containing huts and nesting material for enrichment were changed weekly. Swiss Webster mice were fed standard chow (Teklad 8604 chow) and water *ad libitum.* C57BL/6 mice were fed a diet containing 40 mg/kg doxycycline (DOX) (Bio-Serv F4159).

### 2.2 Stereotaxic Surgery

Aseptic surgeries were performed with mice mounted in a stereotaxic frame (Kopf Instruments) with the skull flat resting on a heating pad. Swiss Webster mice were anesthetized with 5% isoflurane and 80% oxygen (induction), and isoflurane was reduced to 3% afterwards (maintenance). C57BL/6 mice were anesthetized with 4% isoflurane and 80% oxygen (induction), and isoflurane was reduced to 2% afterwards (maintenance). Leading up to surgery, mice were handled for 4 days (3 min per day). Mice were weighed immediately before surgery and given an injection of carprofen (10mg/kg, s.c.). They were also given an injection of saline (0.2ml, s.c.) and ophthalmic ointment was placed over their eyes. The surgical area was prepped by removing the hair with Nair, and then swabbing with 95% isopropyl alcohol solution, betadine, and lidocaine with 2% epinephrine. An incision was made, and the skin was pulled back with two bulldog clamps. Holes were drilled with a microdrill, and viral microinjections were administered via a 10 μL gas-tight Hamilton syringe attached to a microinfusion pump (UMP3, World Precision Instruments), at 100 nL/min. These were targeted bilaterally to the VLO (AP+2.6, ML+/-1.0, DV -2.6 relative to Bregma) (Figure 1A) (Exp. 1) or the dDG (AP-2.2, ML+/-1.3, DV -2.0 relative to Bregma (Exp. 2), and each mouse received 400nl of virus in the VLO (Figure 1A; 4C) or 300nl of virus in the dDG (Figure 4A, D). Post-infusion, the needle was left in place for an additional 10 minutes. Following the injections, the opening was sutured, and the area was reinforced with tissue adhesive. At the end of surgery, mice were weighed again, placed on a heating pad until they were ambulatory, and then placed in a clean cage with hydrogel (mixed with powdered DOX food for Exp. 2), food pellets at the bottom of the cage, and new nesting material. Post-operatively, 16 hours later, mice were weighed and given another injection of carprofen (10mg/kg, i.p.). After another 16 hours, mice were weighed again and given meloxicam (5 mg/kg, s.c.). Apart from cage changes and being weighed, mice were left undisturbed for 7 days to allow for recovery. Experimental procedures did not start until 2 weeks later to allow for viral expression which took 3 weeks to express.

**Figure 1.**
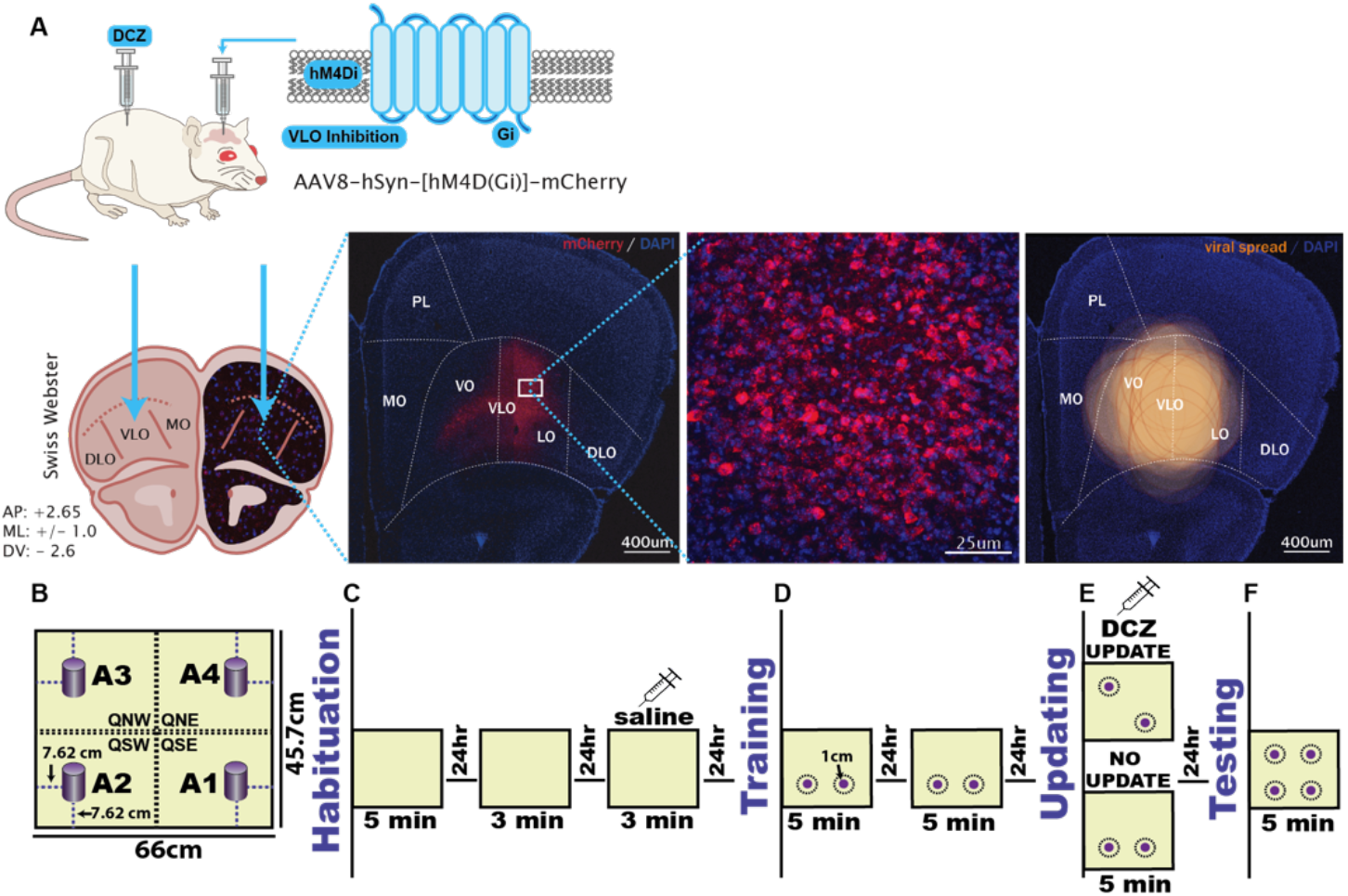
Exp. 1: Viral Strategy and Experimental Design. **A)** Three weeks before behavioral testing, mice received bilateral VLO injections of either an inhibitory DREADD vector (AAV8-hSyn-hM4Di-mCherry) or a control vector (AAV8-hSyn-mCherry). **B–C)** Mice were habituated to a gray chamber for 3 days (5 min on day 1; 3 min on days 2–3). No objects were present, and saline was administered 15 min before the final session. **D)** Mice were then trained for 2 days (5 min/day) with objects in locations A1 and A2. **E)** During the 5-min Updating session, mice were assigned to Update or No-Update groups; the Update group experienced an object-location change (A2→A3). All mice received DCZ (0.1mg/kg, i.p) 15 min beforehand. **F)** On Test day, all four locations (A1–A4) were available: A1 and A2 were familiar; A3 was familiar only to the Update group; A4 was novel. Quadrant time and speed were recorded during Habituation; on all other days, object investigation (within 1 cm) was used to compute Discrimination Indices and Total Exploration.

### 2.3 Viral Microinjections

In Exps. 1 & 2, mice were divided into two groups: experimental mice (hM4Di; Exp. 1 n=20, Exp. 2 n=12) received inhibitory DREADDs AAV_8_-hSyn-hM4D(Gi)-mCherry (Addgene 50475 - titer: 2.8×10^13 GC/mL) while control mice (mCherry; Exp. 1 n=20, Exp. 2 n=12) received AAV_8_-hSyn-mCherry (Addgene 114472 - titer: 2.8×10^13 GC/mL) (Figure 1A). In Exp. 2, to label dDG cells involved in encoding the Updating session, all mice also received bilateral infusions of a viral cocktail consisting of pAAV9-cFos-tTa (UMass Vector Core - titer: 2.7×10^13 GC/mL) and pAAV9-TRE-eYFP (UMass Vector Core - titer: 1.6×10^13 GC/mL).

### 2.4 Drug Preparation

All mice received an injection of deschloroclozapine dihydrochloride (DCZ) 15 minutes prior to the Updating session at a dose of 0.1mg/kg (i.p.). A stock solution of DCZ (HelloBio, HB9126 water soluble) was made at a concentration of 0.02mg/ml with a correction factor of 1.25 to account for the dihydrochloride component of the drug, dissolved in sterile saline (0.9% NaCl). The drug was made on the day of use.

### 2.5 Handling

Mice were handled in a quiet testing room for 3 minutes / day across 4 consecutive days before the start of Habituation. During handling, each mouse was gently grasped and allowed to explore the experimenter’s hand to reduce stress associated with human interaction. On the last day of handling, the mice received saline injections (0.1 ml, i.p.) to habituate them to this aspect of the study.

### 2.6 Apparatus and Data Collection

The experimental apparatus consisted of a grey painted plastic bin (45.7cm W x 66cm L x 30.5cm H) with similarly colored vinyl flooring (Figure 1B). The apparatus was placed on a metal table (88 cm height) in a closed behavioral testing room illuminated with red lighting. The apparatus was open to the room so the mice could use extra maze cues to identify their surroundings and orientation. Fans were turned on to reduce the influence of external environmental sounds. During Training, Updating, and Testing, mice explored identical objects consisting of stainless-steel chrome-plated 500g calibration weights (SEUnMUK, Amazon, 2 inW x 3inH) placed inside plastic containers wrapped in purple lab tape. Each of the four identical objects was placed 7.62 cm from the wall of the apparatus. A top-down camera was mounted directly above the apparatus and connected to a computer system for video recording. Recorded videos were analyzed using EthoVision XT software (Noldus Information Technology, Version 17.5.1718). We recorded the time spent traversing the apparatus and exploring the object-locations (A1-A4) for the various tests (seconds). We also recorded total distance traveled (m) and speed (m/s). For all measures, we used the animal’s centerpoint as a reference point for movement; however, for object investigation we used the mouse’s nosepoint. Object investigation was only included when the mouse’s nose was within 1cm of the object, pointed directly at that object. Between sessions, the apparatus was cleaned with 70% ethanol and wiped down.

### 2.7 Habituation

Habituation took place across 3 consecutive days (Figure 1C). Mice were first habituated to the apparatus for 5 minutes on day 1 (no objects present). On day 2, they were re-exposed for 3 minutes, and on day 3, they underwent a final 3-minute session. Fifteen minutes prior to this session they received a saline injection (0.1 ml, i.p.). Here, we assessed time spent in each quadrant (QSE, QSW, QNW, QNE) to examine whether mice had an inherent preference for any particular area of the apparatus (Figure 2A).

**Figure 2.**
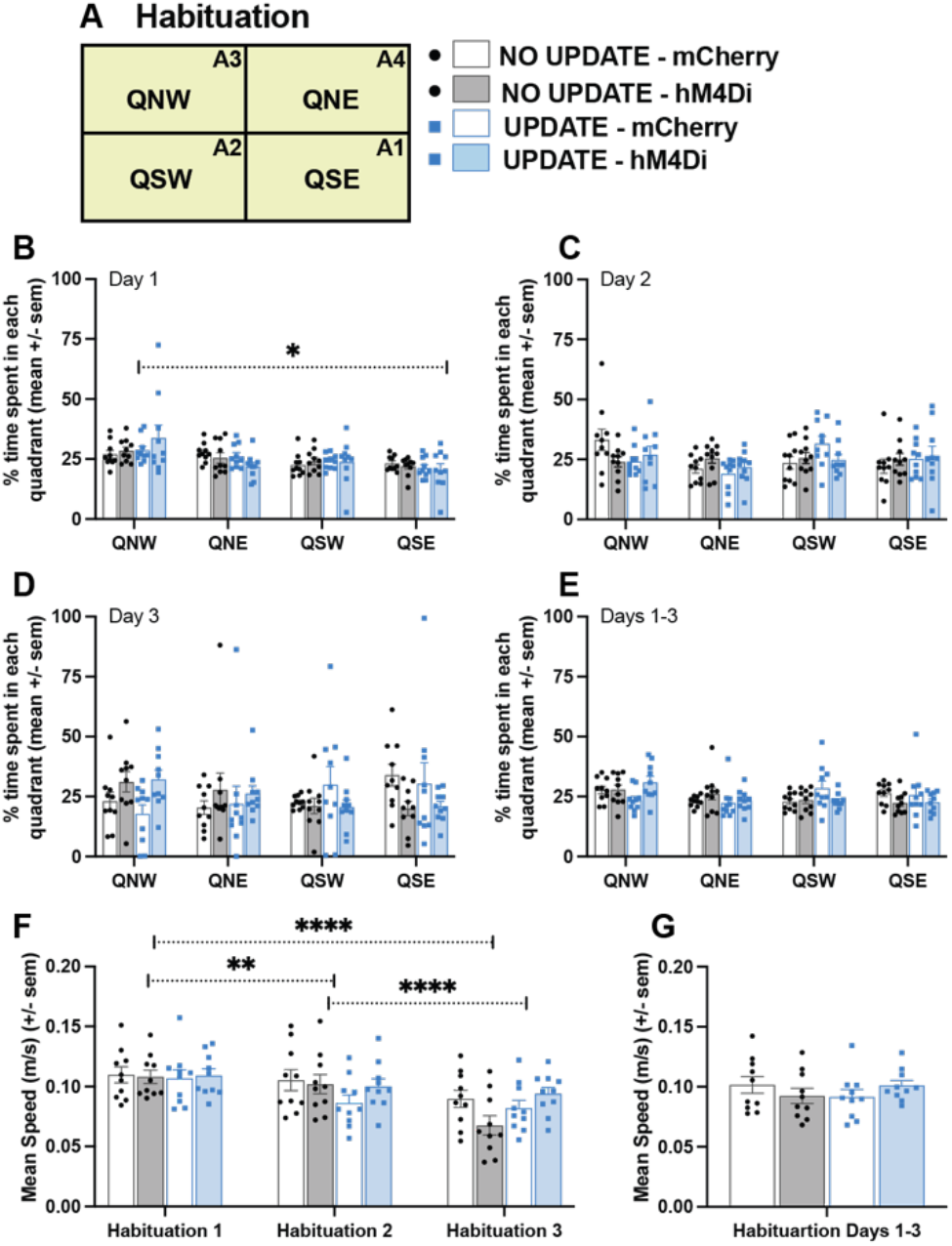
Exp. 1: Habituation. **A)** Schematic of the Habituation arena divided into four quadrants (QNW, QNE, QSW, QSE). Mice belonged to one of four groups based on viral vector (mCherry *vs.* hM4Di) and Updating condition (Update *vs.* No-Update). **B–E)** Percentage of time spent in each quadrant on Habituation Days 1–3 and collapsed across Days 1–3. All groups explored quadrants similarly, with no quadrant biases emerging across days. **F–G)** Mean speed across Habituation sessions and collapsed across Days 1–3. Mean speed decreased across days, but no differences were observed between groups. Together, these data indicate comparable baseline exploration and locomotor activity prior to object Training. *p<0.05; **p<0.01; ***p<0.001; ****p<0.0001.

### 2.8 Training

Following the final day of Habituation, the Training phase of the experiment began which lasted 2 days (Figure 1D). Mice were placed in the apparatus containing two identical objects positioned in the southeast (A1; ZSE) and southwest (A2; ZSW) locations and were allowed to explore freely for 5 minutes (Figure 3A). These object-location configurations remained unchanged throughout Training. A discrimination index (DI) was used to assess each animal’s ability to distinguish between the two object-locations as well as to assess whether they displayed a preference for any object-location. To calculate the DI, the time spent exploring each object-location was used in the following formula: (A2-A1)/(A2+A1)*100. The DI is represented on a scale of 0 (no preference) to +/-100 [preference for one object-location; A1(-), A2(+)]. Exploration was scored when the animal’s nose entered a 1 cm area surrounding the object, defining the zones [ZSE(A1), ZSW(A2), ZNW(A3), ZNE(A4)].

**Figure 3.**
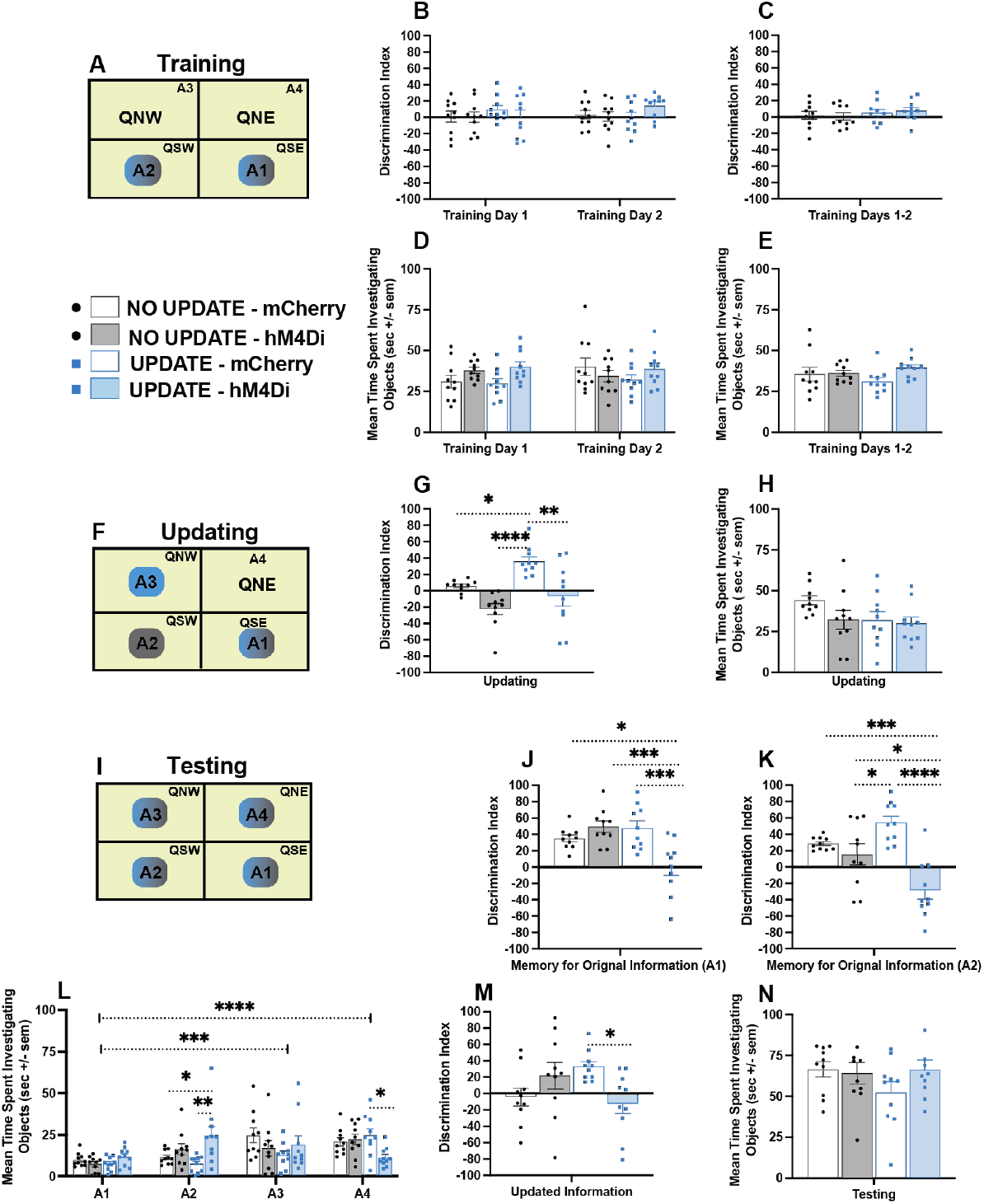
Exp. 1: Training, Updating and Testing. **A)** Schematic of arena during Training showing object-locations A1/A2 were presented to all groups. Mice belonged to one of 4 groups based on virus (mCherry *vs.* hM4Di) and Updating condition (Update *vs.* No-Update). **B)** Discrimination indices (DIs) on Training Days 1–2 and **C)** collapsed across days showing mice explored A1/A2 equally. **D)** Mean time investigating A1/A2 across Training sessions and **E)** collapsed across days. **F)** Schematic of arena during Updating where mice in No-Update groups were given access to A1/A2 again, while mice in the Update groups were given access to A1, but A2 was replaced with object-location A3. **G)** During Updating, control mice preferentially explored the changed location, whereas VLO-inhibited mice failed to do so **H)** Mean time investigating all object-locations during Updating. **I)** Schematic of Testing arena showing all groups were given access to all 4 object-locations including new object-location A4. Memory for the original Training was evaluated by calculating DIs comparing exploration of the novel location (A4) to the original Training locations **J)** (A1) (A4-A1)/(A4+A1)*100, and **K) (**A2) (A4-A2)/(A4+A2)*100. **L)** Mean time investigating each object-location during Testing. **M)** Memory for the updated information was assessed by comparing exploration of the novel (A4) and updated (A3) locations: (A4-A3)/(A4+A3)*100. For No-Update animals, both A3 and A4 were novel locations. **N)** Mean time investigating all object-locations. Control mice distinguished novel from familiar locations, whereas VLO-inhibited mice failed to show this preference when they had experienced an update, indicating impaired memory-updating. *p<0.05; **p<0.01; ***p<0.001; ****p<0.0001.

### 2.9 Updating

Mice subsequently underwent a single Updating session with separate object-location conditions that lasted 5 minutes (Figure 1E). Experimental (hM4Di) and control (mCherry) mice were further divided into two groups: No-Update and Update. Mice in the No Update condition were presented with the same objects in the same locations as Training, A1(ZSE) and A2(ZSW), thus, no update to these locations. In the Update condition, object in location A1 was still present, just as in as Training, however, object in location A2 was moved (updated) to a new location A3 (ZNW) (Figure 3F). In the OUL task, preferential exploration of the changed object-location configuration is interpreted as evidence that the animal detected the spatial change relative to its previously acquired representation. The DI for the No-Update group was calculated as (A2-A1)/(A2+A1)*100, and preference for the novel object-location for the Update group was calculated as (A3-A1)/(A3+A1)*100. In addition to promoting memory-updating, the Updating session provided an opportunity to confirm that animals had acquired the original Training memory (Kwapis et al., 2020). All mice received DCZ (0.1 mg/kg, i.p.) 15 minutes prior to the Updating session followed by a Test session 24 hours later.

### 2.10 Testing

Mice underwent a Test session to assess memory for both the original Training and the updated information (Figure 1F). During the 5-minute Test, mice were presented with four identical objects: three located in previously experienced positions: A1[ZSE], A2[ZSW], A3[ZNW], and one placed in a novel location: A4[ZNE] (Figure 4A). Memory for the original Training was evaluated by calculating DIs comparing exploration of the novel location (A4) to the original Training locations (A1 and A2): (A4-A1)/(A4+A1)*100, (A4-A2)/(A4+A2)*100. Memory for the updated information was assessed by comparing exploration of the novel (A4) and updated (A3) locations: (A4-A3)/(A4+A3)*100. For No-Update animals, both A3 and A4 were novel locations. Each pairwise comparison was designed to address a distinct experimental question. Comparisons of A4 with A1 and A2 assessed memory for the original Training configuration, whereas comparison of A4 with A3 assessed memory for the updated spatial information. Thus, the discrimination indices were intended to evaluate specific memory representations rather than overall exploration across all four locations.

**Figure 4.**
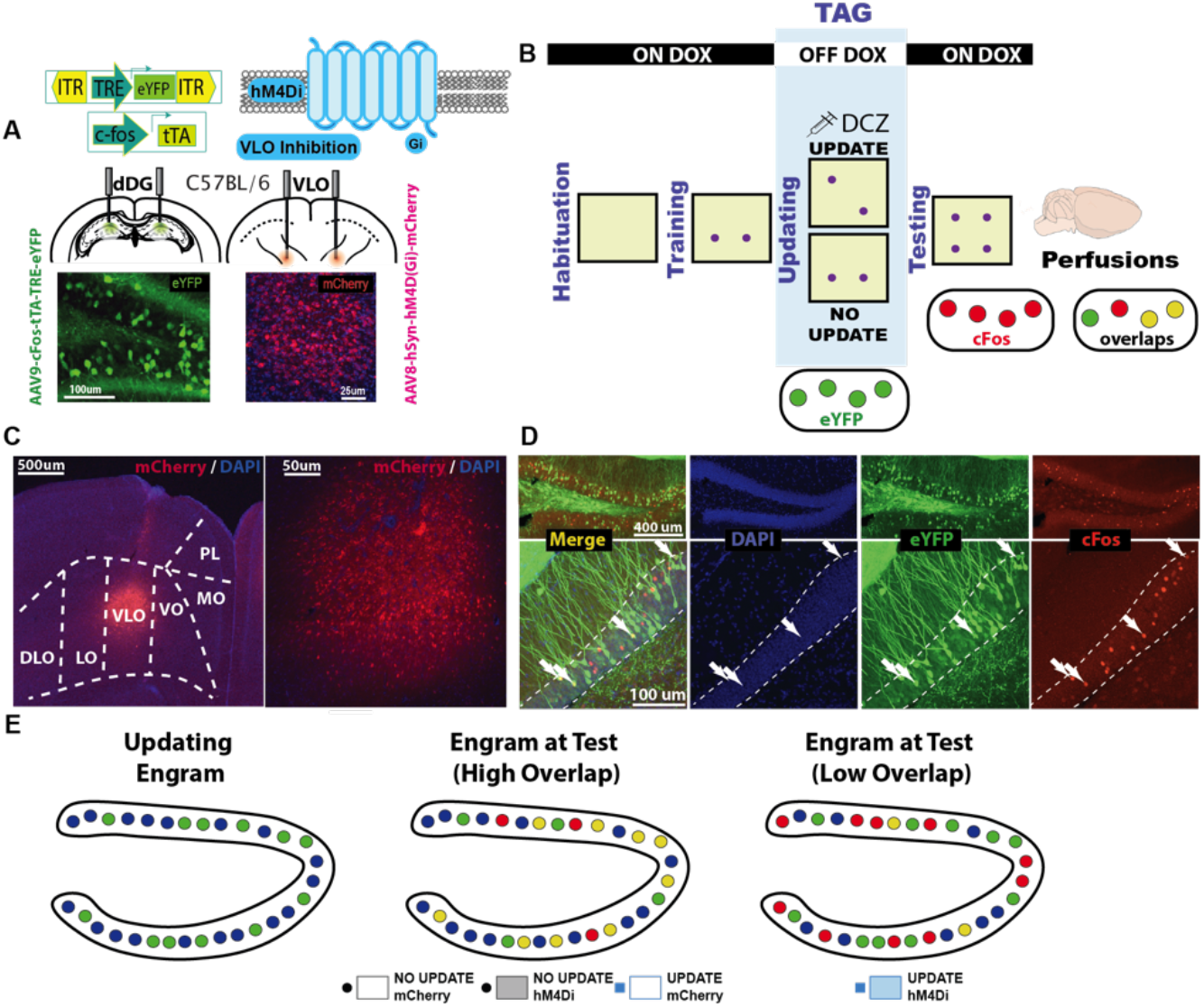
Exp. 2: Effect of VLO Inhibition on Hippocampal Updating-Ensembles. **A)** Viral strategy used: 3 weeks before testing, C57BL/6 mice received bilateral VLO injections of either an inhibitory DREADD vector (AAV8-hSyn-hM4Di-mCherry) or a control vector (AAV8-hSyn-mCherry) as well as bilateral dDG injections of the Tet Tag viral cocktail (AAV9-cFos-ttA-TRE-eYFP). **B)** Mice were habituated to a gray chamber for 3 days (5 min on day 1; 3 min on days 2–3). No objects were present, and saline was administered 15 min before the final session. Mice were then trained for 2 days (5 min/day) with objects in locations A1 and A2. Following the last Training session, mice were taken off DOX to tag the cells encoding the Updating session. Forty-eight hours later, during the 5-min Updating session, mice were assigned to Update or No-Update groups; the Update group experienced an object-location change (A2→A3). All mice received DCZ (0.1mg/kg, i.p.) 15 min beforehand. After the session, the were immediately placed back on DOX to close the tagging window. On Test day, all four locations (A1–A4) were available: A1 and A2 were familiar; A3 was familiar only to the Update group; A4 was novel. Mice were perfused 90 min after the Test session. Quadrant time and speed were recorded during Habituation; on all other days, object investigation (sec, within 1 cm) was used to compute Discrimination Indices and Total Exploration. **C)** Histological representation of viral injection at 5x and 20x in the VLO **D)** and at 10x and 20x in the dDG. Tagged engram in the dDG: DAPI (blue, enclosed within white dotted lines), eYFP (green), cFos (red), overlaps (yellow). **E)** Hypothesized effect of VLO inhibition on dDG ensemble reactivation. Hypothetical tagged engram during Updating session (green) in mCherry and hM4Di groups compared to reactivated engram at Test in cases where the engram at Test includes newly recruited cells (red) and overlapping cells with the Updating session (yellow); DAPI-counterstained cells (blue).

### 2.11 Novelty

On Training day 1, object locations A1 and A2 are both novel. On Training day 2, they both become familiar. During Updating, A1 is familiar to all mice. A2 is presented again to No-Update mice and is also familiar. In Update mice, an object is presented in location A3 instead of A2, violating the animal’s expectations. This object location is familiar. On Test day, all four identical objects are presented in positions A1-A4. A4 is novel for all mice. A3 is novel to No-Update mice and familiar to Update mice. A1 and A2 are familiar to all mice.

### 2.12 Immunohistochemistry

Mice were overdosed with isoflurane and perfused transcardially with (4 °C) phosphate-buffered saline (1x PBS) followed by 4% paraformaldehyde (PFA) in 1x PBS. Brains were extracted and stored overnight in PFA at 4 °C and transferred to a solution of 0.01% sodium azide (S227I-25, Fisher Sci) in 1x PBS the next day. Brains were sectioned into 50 μm coronal slices with a vibratome (Leica, VT100S) and collected in cold 1x PBS. Sections stored in azide were transferred to well plates and washed 3x (10 minutes) in 1x PBS before blocking. They were then blocked for 90 minutes at room temperature in 1x PBS + 2% Triton (PBS-T) and 5% normal goat serum (NGS) on an orbital shaker. Sections were then placed in primary antibody made in PBS-T NGS. To verify VLO injections in Exps. 1 & 2, they were placed in rabbit anti-mCherry (1:1000, Fisher PA5-34974). To quantify overlaps between tagged Updating cells in the dDG (eYFP) and cFos expression in cells at recruited at Test (mCherry), they were placed in chicken anti-GFP (1:1000, Invitrogen a10262) and rabbit anti-cFos (1:1000, Abcam ab190289) respectively. Tissue was then incubated on an orbital shaker at 4 °C for 48 h. Sections were then washed 3x (10 minutes) in PBS-T followed by a 2-hour incubation with secondary antibody Alexa 555 anti-rabbit (1:200, Invitrogen A21428, red) and Alexa 488 anti-chicken (1:200, Invitrogen, A11039, green) made in PBS-T NGS. Following three additional 10-minute washes in PBS-T, sections were mounted onto micro slides (VWR). Nuclei were counterstained with DAPI added to Vectashield HardSet mounting medium (Vector Labs, H1500-10). Slides were then cover slipped and put in the fridge overnight to cure. The following day the edges were sealed with clear nail-polish, and the slides were stored in a slide box in the fridge until imaging.

### 2.13 Fluorescent Confocal Image Acquisition and Quantification

Images were collected from coronal sections using a fluorescent confocal microscope (Zeiss LSM 880) at 5, 10, and 20× magnification with Zen Black software. For histological verification of viral injections and for quantification of overlaps for animals receiving bilateral viral dDG injections, three 20x z-stacks (step size 1μm) were taken from three different slices yielding ∼6 total z-stacks per animal per region. For overlaps, data from each hemisphere was then pooled and the means for the 6 z-stacks were computed. These means were then used to obtain a group mean. Percentage of immunoreactive (eYFP or mCherry) neurons, including overlaps, was defined as a proportion of total DAPI-labeled cells (e.g., eYFP^+^/DAPI*100). Chance overlap was calculated as the percentage of the first immunoreactive neuron multiplied by the percentage of the second immunoreactive neuron (i.e., %eYFP*%cFos). Overlaps were normalized to chance by dividing the proportion of overlaps (i.e., overlaps/DAPI*100) by this chance calculation.

### 2.14 Experimental Design and Statistical Analyses

Across all experiments, we included male and female mice. In Exps. 1 & 2, sex was initially analyzed as an independent variable, but no significant effects or interactions were detected, thus males and females were pooled for all final analyses. Exp. 1 included two primary between-subjects factors: VIRUS condition (hM4Di *vs*. mCherry) and UPDATING condition (Update *vs.* No-Update). We had 20 hM4Di mice and 20 mCherry controls, and each viral group was evenly split into Update and No-Update subgroups (n=10 per subgroup, with 5 males and 5 females in each). Most analyses used three-way repeated-measures (RM) ANOVAs, with Virus and Updating as between-subjects factors and a within-subjects factor such as Quadrant, Zone, or Time (e.g., session). When no RM factor was relevant, for instance, when we examined cell counts and overlaps in Exp. 2, we used two-way ANOVAs with VIRUS and UPDATING as between-subjects factors. For all analyses, the specific statistical test used is clearly stated, and Tukey’s HSD post-hoc tests were applied for follow-up comparisons. Calculated statistics are presented as means ± standard error of the mean (SEM). All statistical tests were conducted in Graphpad Prism (version 10.6.1), were two-tailed, and assumed an alpha level of 0.05. For all figures, *= p<0.05, **= p<0.01, ***= p<0.001, ****= p<0.0001.

## 3. Results

### 3.1 Viral Strategy and Experimental Design

Our virus strategy consisted of targeting the VLO with inhibitory DREADDs in outbred Swiss Webster mice (Figure 1A). Only correctly targeted animals were included in the data analysis (Exp 1: n=40, no excluded mice; Exp 2: n=24, no excluded mice) (Figure 1A). Our paradigm was adapted from Kwapis et al., (2020) who developed the OUL task to assess memory-updating. In this task, mice were exposed to four identical objects in specific locations (Figure 1B). Our design consisted of 3 days of Habituation (Figure 1C) followed by 2 days of Training (Figure 1D). The following day, mice received injections of DCZ 15 minutes prior to the Updating session (Figure 1E) and the next day they were tested for memory retention of the original and updated information (Figure 1F).

### 3.2 Exp. 1: Habituation

During Habituation, mice were placed in the empty chamber with no objects (Figure 2A). We assessed how much time they spent in each quadrant to ensure there was no initial preference across all 3 days of Habituation (Figure 2B-E). On day 1, there was a main effect of QUADRANT (F_3,108_ = 7.268, p=0.0002; 3-way RM ANOVA), an effect driven primarily by two mice in the Update-hM4Di group spending more time in the NW quadrant compared to the SE quadrant (p=0.0172) (Figure 2B). On day 2, there was no effect of QUADRANT (3-way RM ANOVA, *ns*) nor any other group differences (Figure 2C). On the third Habituation day, mice were injected with saline prior to the session. Although they had already been accustomed to receiving injections post-operatively, this was done to habituate them to the injection procedure so that during the updating session they would not elicit a non-specific stress response. During this third Habituation session, we saw more variability in our data set; these injections caused some of the mice to freeze, resulting in them spending more time in one quadrant over the others. We found a QUADRANT x VIRUS interaction (F_3,108_ = 3.549, p=0.0169; 3-way RM ANOVA). However, post-hocs showed no significant differences between any groups (Figure 2D). When we took the mean of the 3 Habituation days collapsed, there were no group differences - all mice spent equal time in each quadrant (3-way RM ANOVA, *ns*) (Figure 2E). We also measured the speed at which the mice traversed the apparatus. Mice traveled slower across the 3 days of Habituation, mainly owing to differences between the third day when they received the saline injection and spent some time immobile (Figure 2F) and the other two days where they moved around more. There was a DAY x VIRUS x UPDATING interaction (F_2,72_ = 3.351, p=0.0406; 3-way RM ANOVA). On day 3, mice traversed the apparatus at a slower velocity compared to day 2 (p<0.0001) and day 1(p<0.0001) and on day 2 they traveled slower than day 1 (p=0.0028). When looking at the mean of the 3 days, there were no significant differences across groups in terms of speed (2-way ANOVA, *ns*) (Figure 2G).

### 3.3 Exp. 1: Training

Training consisted of two sessions where mice in all groups were given access to object-locations A1 and A2 (Figure 3A). During both sessions, mice exhibited a DI close to zero indicating that they explored each object-location equally (Figures 3B-E). We also examined the total amount of time spent exploring, and this was also equal across groups (Figures 3D-E).

### 3.4 Exp. 1: Updating

During the Updating session, mice in the No-Update groups were again given access to object-locations A1 and A2, while mice in the Update groups were given access to A1, but A2 was replaced with object-location A3 instead, reflecting an *updated* event with new information (Figure 3F). Fifteen minutes prior to the session, all mice were injected with 0.1mg/kg DCZ to inhibit the VLO. No-Update control (mCherry) mice explored object locations A1 and A2 equally, consistent with the preceding two Training days (Figure 3G). VLO inhibition in the No-Update condition produced a slight shift toward greater exploration of A2 relative to A1. However, this difference between the No-Update mCherry and No-Update hM4Di groups was not significant.. In the Update condition, control mice preferentially explored the updated location A3, whereas VLO-inhibited mice explored A1 and A3 equally. This was clear as we saw a main effect of UPDATING (F_1,36_ = 8.695, p=0.0056; 2-way ANOVA). Mice in the mCherry Update control group had a higher DI compared to mCherry controls in the No-Update condition (p=0.0456). When new information was presented in the Update condition, VLO inhibition resulted in mice spending equal time in both A1 and A3 object locations. Thus, VLO inhibition abolished the normal preference for the updated object-location configuration observed in controls. We also saw a main effect of VIRUS (F_1,36_ = 20.53, p<0.0001; 2-way ANOVA) where overall, VLO inhibition biased mice towards interacting with object location A1. Importantly, these group differences cannot be attributed to overall exploration time, which did not differ across groups (Figure 3H).

### 3.5 Exp. 1: Testing

#### 3.5.1 Memory for Original Information

During the Testing session, mice in all groups were given access to all four object-locations including new object-location A4 (Figure 3I). Memory for familiar / original information, object-locations A1 and A2 were assessed by comparing time spent in object-location A4 to A1 and A2. When looking at a comparison of A4 to A1 (Figure 3J) we saw a VIRUS x UPDATING interaction (F_1,36_ = 15.95, p=0.0003; 2-way ANOVA). When looking at a comparison of A4 to A2 (Figure 3K) we also saw a VIRUS x UPDATING interaction (F_1,36_ = 13.34, p=0.0008; 2-way ANOVA). Mice in both the No-Update groups (mCherry and hM4Di) spent more time investigating object-location A4 compared to both familiar locations A1 (Figure 3J) and A2 (Figure 3K), evidenced by a positive DI. In the No-Update condition, VLO inhibition the previous day did not have an effect (Figure 3J: p=0.05611; Figure 3K: p=0.7595). Similarly, mice in the Update-mCherry group preferentially explored A4 over both familiar object-location configurations, A1 (Figure 3J) and A2 (Figure 3K). However, VLO inhibition the prior day (during the presence of new object-location A3), resulted in VLO-inhibited mice in the Update condition spending less time with object-location A4 and more time with familiar object-locations A1 (Figure 3J: p=0.0006) and A2 (Figure 3K: p<0.0001) compared to Update-mCherry mice. In Figure 3K, mice in the No-Update-hM4Di group were also significantly different than Update-mCherry controls suggesting that inhibitory DREADDs in general, were biasing the mice towards object-location A2 (p=0.031). During the Testing session, we also looked at time spent at each object-location across groups and mice in the Update-hM4Di condition spent more time at location A2: main effect of VIRUS (F_1,36_ = 11.13, p=0.002; 2-way ANOVA) compared to controls (no Update mCherry p=0.0411; Update mCherry p=0.0043) (Figure 3L). Mice in the Update-hM4Di group also spent less time at object-location A4 compared to other groups (p=0.0126). Overall, all mice spent more time spent in location A3 (p = 0.0006) and A4 (p < 0.0001) compared to A1 during this session: OBJECT x VIRUS x UPDATING interaction (F_3,108_ = 4.225, p=0.0072; 3-way RM ANOVA).

#### 3.5.2 Memory for Updated Information

Memory for updated information in position A3 was assessed by comparing time spent at locations A4 to A3. When looking at a comparison of A4 to A3 (Figure 3M) we again see a VIRUS x UPDATING interaction (F_1,36_ = 9.183, p=0.0045; 2-way ANOVA). Both No-Update groups (mCherry and hM4Di) spent comparable time exploring A3 and A4, indicating no preference between the two novel object-location configurations. VLO inhibition did not alter this pattern of exploration. Similarly, Update-mCherry mice spent more time exploring object-location A4 than A3, demonstrating a preference for the novel over the updated object-location configuration. However, update-mCherry spent the least amount of time at object locations A1 (p=0.0248) and A2 (p=0.0218) compared to A4, which were even more familiar than A3 (Figure 3L). VLO inhibition during the Updating session resulted in Update-hM4Di mice spending equal time exploring object locations A3 and A4 at Test, whereas Update-mCherry mice preferentially explored A4 (p=0.0478). This was not due to differences in the total amount of time spent investigating objects as this was equal across groups (Figure 3N).

### 3.7 Exp. 2: Effect of VLO Inhibition on Hippocampal Updating-Ensembles

Although Exp. 1 demonstrated that VLO inhibition during the Updating session impaired performance in the OUL task during both Updating and Test, the neural basis of this behavioral impairment remained unknown. We therefore performed a second experiment (Exp. 2) to determine whether VLO inhibition altered hippocampal ensemble dynamics associated with the Updating experience. To accomplish this, we repeated the behavioral experiment using C57BL/6J mice, a strain compatible with the TetTag activity-dependent tagging approach, which also allowed us to determine whether the behavioral findings generalized across mouse strains. The behavioral procedures were identical to those used in Exp. 1. Neurons recruited in the dDG during the Updating session were tagged (Figure 4A) and later compared with neurons expressing cFos during the Test session (Figure 4B). This approach allowed us to quantify reactivation of the Updating-tagged dDG ensemble and determine whether VLO inhibition (Figure 4C) altered subsequent hippocampal ensemble reactivation (Figures 4D–E).

#### 3.7.1 Behavior

On the first day of Habituation (Figure 5A), there was a main effect of QUADRANT (F_3,60_ = 3.310, p=0.026; 3-way RM ANOVA), an effect driven primarily by all mice spending slightly more time in the NW quadrant compared to the NE quadrant (p=0.0227) (Figure 5B). Following that, there were no other group differences during Habituation, and all mice spent equal time across quadrants (3-way RM ANOVA, *ns*) (Figure 5C-E). Mice did traverse the maze quicker on day 1 compared to the other two days (Figure 5F). There was a main effect of DAY (F_2,40_ = 55.99, p<0.0001; 3-way RM ANOVA) with a significant difference between day 1 and day 2 (p < 0.0001), day 1 and day 3 (p<0.0001) and day 2 and day 3 (p=0.0079) with mice getting progressively slower. However, there were no group differences.

**Figure 5.**
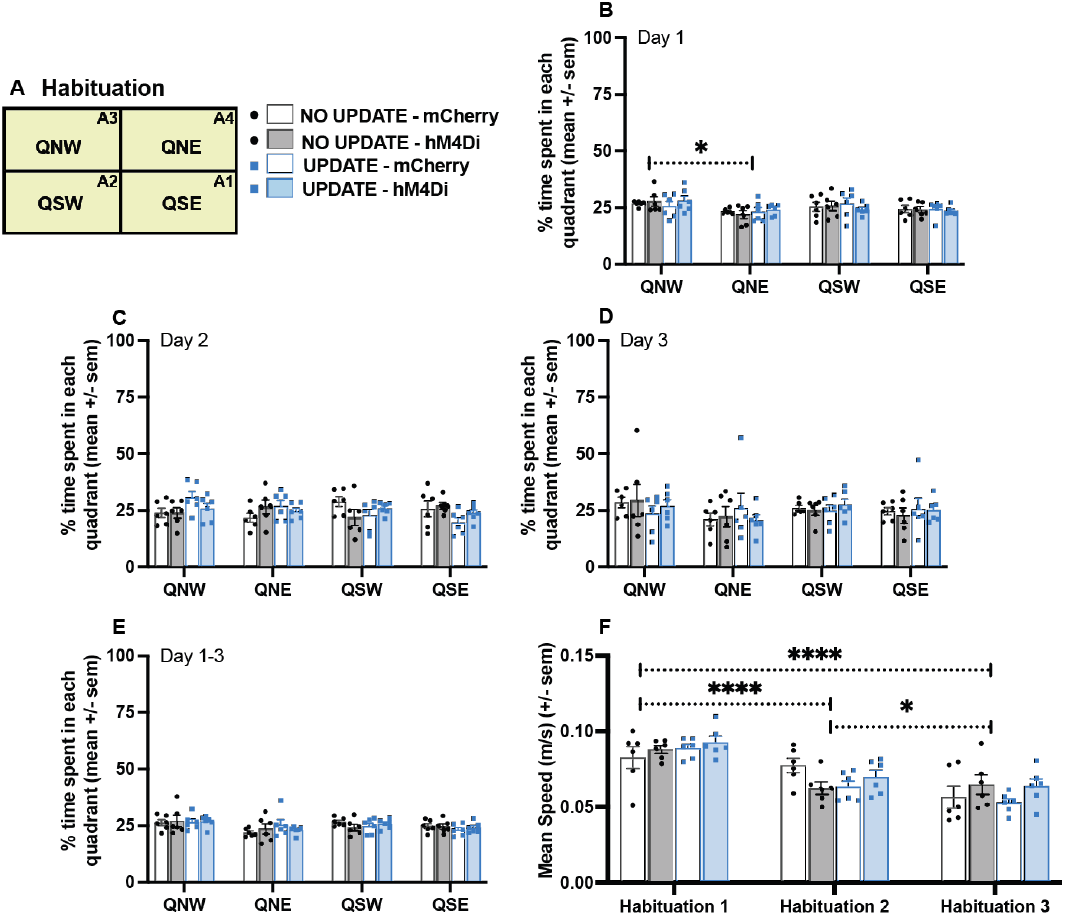
Exp. 2: Habituation. **A)** Schematic of the Habituation arena divided into four quadrants (QNW, QNE, QSW, QSE). Mice belonged to one of four groups based on viral vector (mCherry *vs.* hM4Di) and Updating condition (Update *vs.* No-Update). **B–E)** Percentage of time spent in each quadrant on Habituation Days 1–3 and collapsed across Days 1–3. All groups explored quadrants similarly, with no quadrant biases emerging across days. **F)** Mean speed across Habituation sessions. Mean speed decreased across days, but no group differences were observed between groups indicating comparable baseline exploration and locomotor activity prior to object Training. *p<0.05; **p<0.01; ***p<0.001; ****p<0.0001.

During Training (Figure 6A) mice exhibited a DI close to zero indicating that they explored each object-location equally (Figures 6B-C). All mice spent more time exploring on Training day 1 compared to day 2 (Figure 6D). There was a main effect of DAY (F_1,20_ = 28.97, p<0.0001; 3-way RM ANOVA) with a significant difference across days in all groups except the Update hM4Di group (No-Update mCherry: p=0.0094; No-Update hM4Di: p=0.0473; Update mCherry: p=0.0345).

**Figure 6.**
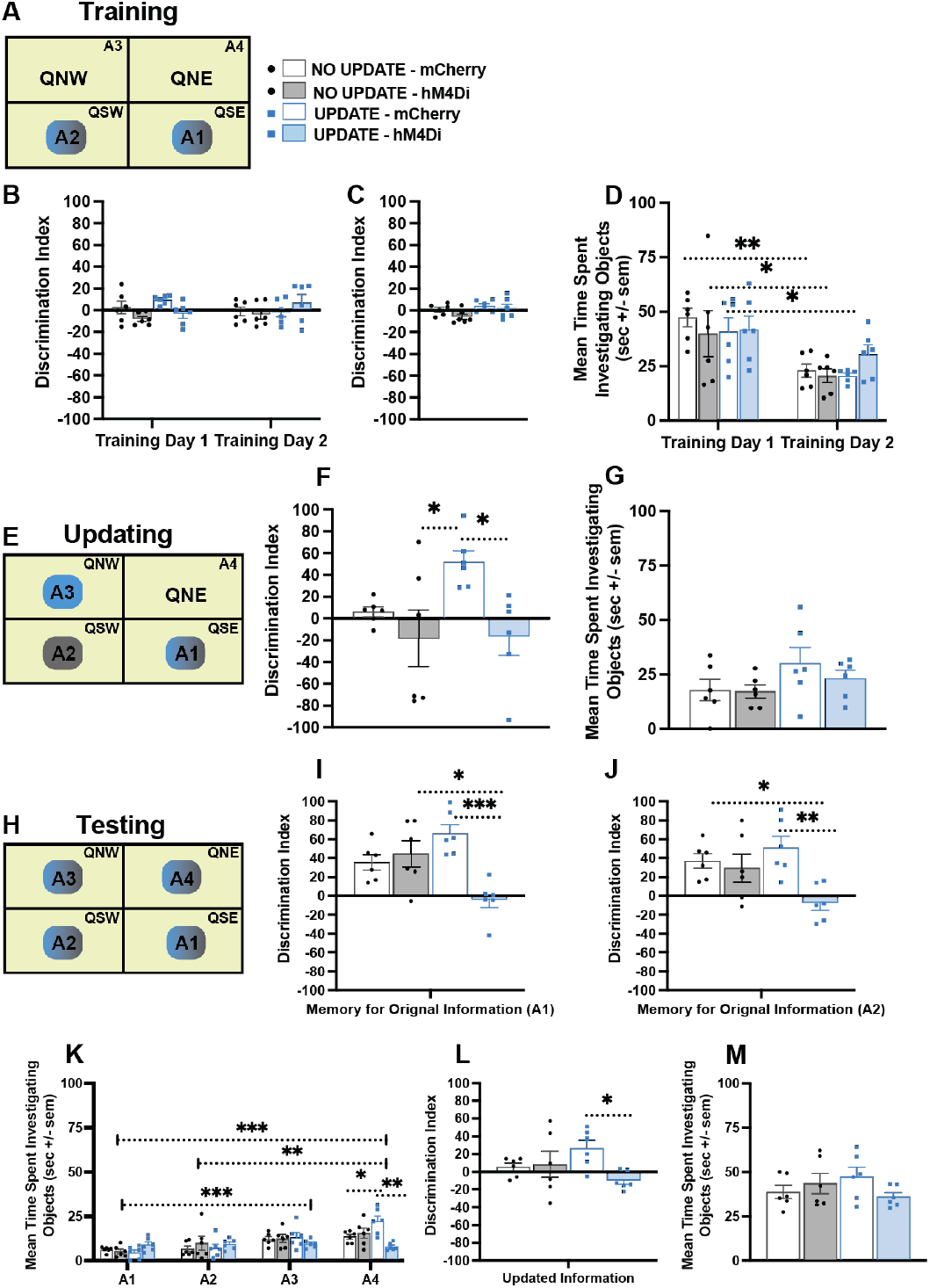
Exp. 2: Training, Updating and Test. **A)** Schematic of arena during Training showing that object-locations A1/A2 were presented to all groups. **B)** Discrimination indices (DIs) on Training Days 1–2 and **C)** collapsed across both days **D)** Mean time spent investigating objects in locations A1 and A2 across Training sessions and **E)** Schematic of arena during Updating session. During Updating, No-Update mice revisited object-locations A1 and A2, while Update mice revisited A1, but saw A3 instead of A2. **F)** DIs during Updating: control mice preferentially explored the changed location, whereas VLO-inhibited mice failed to do so **G)** Mean time spent investigating all object-locations during Updating session. **H)** Schematic of Testing arena showing all groups were given access to all four object-locations including new object-location A4. **I-J)** DIs showing memory for original information. **K)** Mean time spent investigating each object-location during the Testing session. **L)** DIs showing knowledge of the updated information. **M)** Mean time spent investigating all object-locations. Similar to Exp. 1, control mice distinguished novel from familiar locations, whereas VLO-inhibited mice failed to show this preference when they had experienced an update, indicating impaired memory-updating. *p<0.05; **p<0.01; ***p<0.001; ****p<0.0001.

During the Updating session, No-Update mice revisited object-locations A1 and A2 again, while Update mice revisited A1, but were given an object in location A3 instead of A2 (Figure 6E). As before, mice received DCZ 15 minutes before the session to silence VLO activity. No-Update control (mCherry) mice explored A1 and A2 equally (Figure 6F). VLO inhibition in the No-Update condition produced a slight shift toward greater exploration of A2 relative to A1; however, this difference between the No-Update mCherry and No-Update hM4Di groups was not significant. In the Update condition, mCherry mice treated object-location A3 as novel, exploring here more than object-location A1 (Figure 6F). There was a main effect of VIRUS (F_1,20_ = 7.885, p=0.0109; 2-way ANOVA). VLO inhibition in the presence of new information (A3) impaired mice as before, causing the animals to spend more time with objects in familiar location A1 compared to A3. Update-mCherry mice differed from both Update-hM4Di (p=0.0383) and No-Update hM4Di mice (p=0.0331). This deficit was not related to overall exploration time, as it did not differ across groups (Figure 6G).

During Testing, mice were given access to all four object-locations including new object-location A4 (Figure 6H). Memory for Original Information: Comparing time spent in location A4 to A1, there was a VIRUS x UPDATING interaction (F_1,20_ = 15.18, p=0.0009; 2-way ANOVA). All mice including the No-Update hM4Di mice explored object-location A4 more than A1. However, Update-hM4Di were impaired in this respect (Figure 6I). With a DI close to zero, they explored both object-locations equally. DIs were significantly different between No-Update hM4Di mice and Update-mCherry mice (p=0.0005), as well as No-Update hM4Di mice (p = 0.0143). When comparing time spent in object location A4 to A2 (Figure 6J), we also saw a VIRUS x UPDATING interaction (F_1,20_ = 5.32, p=0.0319; 2-way ANOVA). Both No-Update groups (mCherry and hM4Di) preferentially explored A4 over A1, indicating that VLO inhibition during the preceding Updating session did not alter this pattern of exploration. Similarly, mice in the Update control condition also preferentially explored object-location A4 as novel in comparison to A1. As we saw in Exp. 1, VLO inhibition the prior day (during the presence of new object-location A3), resulted in Update-hM4Di mice spending less time exploring object-location A4 and more time exploring object-location A1. The DI for this group was significantly different compared to controls - No-Update mCherry (p=0.0458) and Update mCherry (p = 0.0065). Figure 6K shows that all mice spent more time exploring object-location A4 compared to A1-A3 except the Update-hM4Di mice, which spent equal time exploring all object-locations. There was an OBJECT x VIRUS x UPDATING interaction (F_3,60_ = 5.985, p=0.0012; 3-way RM ANOVA). Memory for Updated Information: Memory for updated information A3 was assessed by comparing time spent at location A4 to A3 (Figure 6L). We again see a VIRUS x UPDATING interaction (F_1,20_ = 4.723, p=0.0419; 2-way ANOVA). Both No-Update groups (mCherry and hM4Di) spent equal time exploring A3 and A4, indicating no preference between these two object-location configurations. VLO inhibition during the preceding Updating session did not alter this pattern of exploration. Update-mCherry mice also preferentially explored object-location A4 compared to A3 while update-hM4Di mice spent equal time exploring both A3 and A4 (p=0.0462). This was not due to differences in the total amount of time spent investigating objects as this was equal across groups (Figure 6M).

#### 3.7.2 Overlaps

To assess whether DREADD-induced VLO inhibition altered reactivation of Updating-related hippocampal ensembles, we quantified the overlap (yellow) between neurons tagged during the Updating session (eYFP⁺, green) and neurons active during the Test session (cFos⁺, red) (Figure 7A). We first confirmed that the total number of DAPI-labeled cells (blue) did not differ across groups (2-way ANOVA, *ns*) (Figure 7B). Likewise, neither the size of the Updating-tagged ensemble, expressed as a percentage of DAPI-labeled neurons (Figure 7C), nor the proportion of cFos⁺ neurons active during Test (Figure 7D) differed between groups (2-way ANOVA, *ns*), averaging approximately 10% and 2.5%, respectively. We then quantified the overlap between these two neuronal populations (Figures 7D–E), which reflects the extent to which neurons recruited during the Updating session were reactivated during Test.

**Figure 7.**
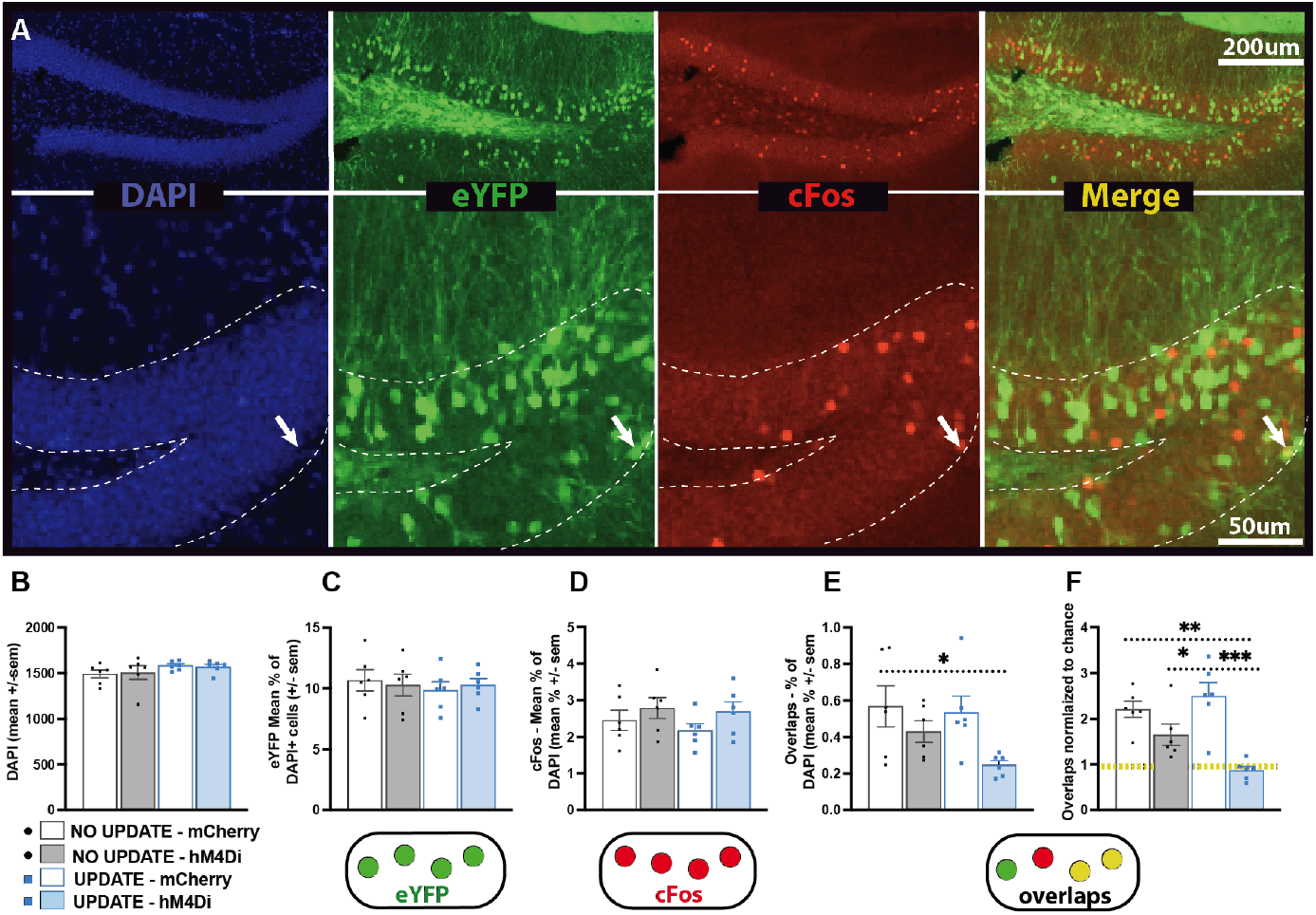
Exp. 2: Overlaps. **A)** Representative dDG images (10x top, 20× bottom) of a tagged engram at Updating (eYFP, green) and the engram recruited during the Test session (cFos, red). Overlap between these engrams can be seen in yellow (merge); DAPI (blue) was used as a counterstain. **B)** Across groups, a similar number of cells were stained with DAPI. **C)** Engram sizes did not differ across groups during Updating **D)** or during Test. **E)** There were fewer overlaps in the experimental VLO-inhibited Update-hM4Di mice, and this matched the behavioral impairment seen in these animals. **F)** This was normalized to chance (yellow dotted line). *p<0.05; **p<0.01; ***p<0.001; ****p<0.0001.

In the No-Update condition, we expected comparable overlap between the mCherry and hM4Di groups if VLO inhibition did not alter reactivation of the Updating-tagged dDG ensemble (Figure 4E). In the Update condition, we hypothesized that VLO inhibition during Updating would reduce subsequent reactivation of the Updating-tagged ensemble, resulting in lower overlap in Update-hM4Di mice relative to Update-mCherry controls. This prediction was based on the hypothesis that VLO activity contributes to establishing or stabilizing hippocampal representations associated with the Updating experience. Consistent with this hypothesis, we observed a main effect of VIRUS (F1,20 = 7.139, p=0.0147; 2-way ANOVA) driven by reduced overlaps in the Update-hM4Di group (p=0.0436) (Figure 7E). When overlaps were normalized to chance (chance = 1) there was a significant VIRUS x UPDATING interaction (F1_,20_ = 6.429, p=0.0197; 2-way ANOVA) (Figure 7F). Overlaps in the Update-hM4Di group were significantly reduced relative to both mCherry control groups (Update: p=0.0001 ; No-Update: p=0.0012). Although VLO inhibition produced only modest effects on DIs in the No-Update condition (Figures 3G, 6E), it also produced a small reduction in normalized overlap in this group (p=0.0463; Figure 7F), suggesting that VLO inhibition may exert subtle effects on hippocampal ensemble dynamics even when no updating demand is present.

## 4. Discussion

We showed that VLO inhibition using inhibitory DREADDs disrupted performance on the OUL task during conditions requiring memory-updating, without impairing novelty exploration or memory for original spatial information. Rodents often demonstrate a preference for novelty (Berlyne, 1950; Clinton et al., 2007; Ennaceur & Delacour, 1988; Lueptow, 2017). Consistent with this pattern of exploratory behavior, mice spend more time exploring novel *vs.* familiar stimuli, a principle used in the OUL paradigm (Kwapis et al., 2020) and other tests e.g., novel object recognition (Lueptow, 2017), social preference (Beery & Shambaugh, 2021; Choleris et al., 2009; Ervin et al., 2015), spontaneous alternation (d’Isa et al., 2021), and the hole board (File & Wardill, 1975). However, novelty preference can vary with strain, age, and environmental enrichment (Arakawa, 2023; Dickson & Mittleman, 2021; Murai et al., 2007) and may reflect multiple processes including exploratory motivation and curiosity, (Swiercz et al., 2025; van Lieshout et al., 2018; M. Z. Wang & Hayden, 2021), which have been tied to the OFC (Blanchard et al., 2015). The disruption we observed is not readily explained by a generalized reduction in novelty-driven exploration. VLO inhibition did not alter novelty preference when no updating demand was present, and deficits emerged specifically when animals encountered new information that needed to be integrated with an existing spatial memory. Thus, the behavioral effects of VLO inhibition emerged specifically when novel spatial information had to be evaluated in the context of an existing memory representation. This dissociation suggests that VLO contributes to evaluating novel information in the context of existing memory representations to support updating, rather than broadly regulating novelty exploration itself. Future studies designed to dissociate exploratory motivation from memory-updating will further clarify the contribution of OFC circuits to these processes.

Behavioral control measures did not account for these deficits. Baseline exploration during Habituation was comparable across groups, with no consistent quadrant preferences or locomotor differences. Although acute saline injections increased variability on the final Habituation day, these effects did not differ between groups, indicating that neither handling nor injection procedures introduced systematic pre-testing differences. Overall, locomotion also did not differ across groups, suggesting subsequent memory-updating impairments were not due to motor deficits or altered baseline exploration.

Training performance confirmed all mice acquired the initial spatial representation, reflected by equal exploration across object-locations. The Updating session further underscored the specificity of the deficit: control mice reliably detected the changed location and preferentially explored it, whereas VLO-inhibited mice failed to treat the updated location as meaningfully different despite equivalent exploration time. This impairment emerged only when new information was introduced; VLO inhibition had little effect on behavior when no update was required. These findings suggest that VLO activity is not required for exploration of novel spatial configurations in the absence of updating demands.

During Updating, mCherry control mice reliably detected the location change and selectively explored the novel A3 position, demonstrating intact prediction error–driven updating. In contrast, VLO-inhibited mice failed to show this novelty preference and instead explored the familiar and updated locations equally, despite normal exploration time. This selective disruption during the presentation of new information absent in the No-Update condition, indicates VLO activity is required to detect or assign salience to unexpected (spatial) changes, rather than supporting general exploration, novelty seeking, or memory for the original configuration. VLO inhibition did not disrupt retrieval or novelty preference in the absence of updating demands. This update-specific deficit aligns with the idea that the VLO contributes to interpreting novelty as informative, that is, detecting when an unexpected change signals a need to revise an existing representation. VLO activity appears to be required when unexpected information must be evaluated against prior expectations.

These findings are consistent with prior work positioning the OFC (Schoenbaum et al., 2011), and particularly the ventrolateral subdivision (Sarlitto et al., 2018), as a hub for encoding outcome expectancies, detecting violations of those expectancies, and updating internal models when predictions change (Chan et al., 2021; Ma et al., 2025; Wilson et al., 2014). Across species, studies show the OFC signals unexpected outcomes, integrates multisensory and associative information, and supports flexible behavioral adjustments when contingencies shift (Riceberg & Shapiro, 2017; B. A. Wang et al., 2023). Our findings extend this role beyond reward contexts by demonstrating VLO activity is similarly required when change is expressed in a spatial memory domain. Where prior studies have emphasized reversal learning and value-updating (Amodeo et al., 2017; Rudebeck & Murray, 2014), the present work shows the same computations support hippocampal-dependent memory-updating. Thus, the dissociation we observed, intact novelty preference when novelty did not challenge an existing memory, but impaired novelty-guided behavior when novelty required comparison to prior knowledge, implicates the VLO in evaluating change rather than simply detecting it, situating the VLO within a predictive updating framework.

At Test, memory for the original spatial configuration remained intact across all groups, except specifically when the VLO was inhibited during Updating. Both control groups and the No-Update hM4Di group identified A4 as novel, demonstrating reliable memory for A1/A2. However, VLO-inhibited mice failed to show this novelty preference at Test and instead shifted their exploration toward familiar locations. This selective disruption indicates that suppressing VLO activity at the time new information was presented interfered with stability or accessibility of the original memory trace, or altered the weighting of familiarity signals, without affecting baseline novelty detection or retrieval processes. This pattern is consistent with the OFC’s established role in inferring and updating mental representations of *What situation am I in now?* (Wilson et al., 2014). When VLO activity was suppressed during the prediction error moment, the internal model guiding familiarity and novelty judgments appeared to be mis-updated or incompletely updated, since the VLO was offline, leading to distorted representations of what was familiar, updated, or novel at Test.

A similar pattern emerged for memory of the updated information. Control mice that experienced an update discriminated A4 from A3 as expected, treating A4 as novel while retaining memory of A3 from the previous day. In contrast, VLO-inhibited mice in the Update condition failed to show this distinction and explored A3/A4 equally, indicating an inability to encode or retain the updated spatial representation. Because total exploration time was normal, this deficit reflects a specific failure of memory-updating rather than reduced engagement or impaired novelty detection. The pattern of behavior also suggests a form of perseveration driven by inadequate suppression of prior spatial representations (Morton & Munakata, 2002). Under normal conditions, the OFC, and particularly the VLO, helps shift behavior away from outdated associations once new information becomes relevant (Cerpa et al., 2023; Hervig et al., 2020; Izquierdo et al., 2017; Schoenbaum et al., 2000). When the VLO was inhibited during Updating, mice not only failed to incorporate the new spatial information but also showed an inappropriate persistence of responding to previously familiar locations at Test. This indicates that updating deficits may reflect a dual failure: reduced incorporation of new, prediction error–driven information (Takahashi et al., 2009; Zimmermann et al., 2018) and a parallel inability to de-prioritize or suppress outdated memory traces (Izquierdo et al., 2017; Morton & Munakata, 2002). Such perseverative tendencies are consistent with OFC dysfunction more broadly and align with its established role in inhibiting obsolete associations during flexible behavior (Chudasama & Robbins, 2003).

The behavioral and cellular findings were broadly consistent, indicating VLO activity during Updating supports both accurate memory-guided performance and appropriate hippocampal ensemble reorganization. In the Update-condition, VLO inhibition impaired discrimination behaviorally while also reducing overlap between Updating-tagged and Test-reactivated dDG ensembles, suggesting that disrupted incorporation of new information at the neural level contributes to the observed behavioral deficit. By contrast, control mice maintained both intact performance and higher overlaps, consistent with successful integration of prior and newly acquired representations. Notably, in the No-Update condition, VLO inhibition produced only modest behavioral effects despite some reduction in overlap, implying subtle alterations in ensemble dynamics may be tolerated when task demands do not require active updating.

Future studies should explore underlying neural circuitry of the observed behavioral and cellular effects, specifically OFC-hippocampus pathways. Previous studies show connectivity between the hippocampus and PFC (Sigurdsson & Duvarci, 2016), but direct OFC-hippocampus projections remain poorly characterized (Giarrocco & Averbeck, 2023). OFC projects to the lateral portion of the entorhinal and rostral portion of the perirhinal cortex (Kondo & Witter, 2014; Witter et al., 2017), regions involved in spatial / object recognition (Chao et al., 2022; Connor & Knierim, 2017; Huang et al., 2023). Thus, rather than direct monosynaptic OFC-hippocampus connections, these structures likely communicate through intermediate regions (e.g., entorhinal, perirhinal, amygdala, mediodorsal thalamus) (Murray & Wise, 2010; Rempel-Clower, 2007; Rolls et al., 2020; Zikopoulos & Barbas, 2006). Functionally however, they appear to form a tightly coupled network supporting state representation, model-based learning, and memory-guided decision making. Moreover, we focused our investigation on the dDG because of its established role in pattern separation, memory-updating, and engram formation. However, other hippocampal subregions also contribute to these processes and should be explored in future studies. Thus, an important future direction should be to determine whether VLO inhibition selectively alters ensemble dynamics in the dDG or whether comparable effects are observed in other hippocampal or extrahippocampal regions not directly implicated in memory-updating.

In conclusion, our findings support a critical role of the VLO in updating object-location memories. Our results support a role for the VLO in integrating unexpected information into existing spatial memory representations. Consistent with this, inhibiting VLO activity altered contextual hippocampal engram overlap and caused mice to fail to preferentially explore the updated configuration, leading to a failure to prioritize novel over familiar object-locations. Together, these findings support a role for the VLO in guiding adaptive memory-updating by linking salience detection to reorganization of memory representations that support flexible behavior. Our findings align with a growing body of work across rodents, nonhuman primates, and humans demonstrating that the OFC is critical for updating outcome expectations when new information becomes available (Aguirre et al., 2024; Cerpa et al., 2023; Chudasama & Robbins, 2003; Schoenbaum et al., 2000). These results are consistent with contemporary accounts proposing that the OFC supports inference-based updating and/or credit assignment (Ma et al., 2025; Qiu et al., 2024; Schiereck et al., 2026; Witkowski et al., 2025), while extending these frameworks by demonstrating that VLO is required for updating hippocampal-dependent spatial memories outside reward contexts (Costa et al., 2014; Wilson et al., 2014) and that this process is associated with hippocampal engram reorganization.

## Acknowledgements

We’d like to thank Beata Czesny, Joseph Schluep, and the Animal Care Facility for technical assistance with the project.

## Author Contributions

Each author must be identified with at least one of the following: Designed research (SLG, KK, DY); Performed research (KK, DY, WFW, RZW, ZA, MC, GR, AD, SLG); Analyzed data (SLG); Wrote the first draft of the paper (KK, DY, WFW, RZW, SLG); Edited the paper (KK, DY, WFW, RZW, ZA, MC, GR, AD, SLG).

## Clinical trial

not applicable.

